# Evolutionary dynamics of an epigenetic switch in a fluctuating environment

**DOI:** 10.1101/072199

**Authors:** Mariana Gómez-Schiavon, Nicolas E. Buchler

## Abstract

Adaptation and survival in fluctuating environments is an evolutionary challenge faced by organisms. Epigenetic switches (bistable, molecular systems built from self-reinforcing feedback loops) have been suggested as a mechanism of bet-hedging and adaptation to fluctuating environments. These epigenetic systems are capable of spontaneously switching between phenotypes in the absence of DNA mutation and these phenotypes are stably inherited through multiple cell generations. The extent to which epigenetic switches outcompete adaptation through genetic mutation in fluctuating environments remains unexplored. To better understand the conditions that select for epigenetic switching, we used computer simulation to evolve a mechanistic model of a self-activating genetic circuit, which can adapt genetically and exhibit epigenetic switching. We evolved this model in a fluctuating environment under different selection pressures, mutation step-sizes, population sizes, and fluctuation frequencies. There was a trade-off between minimizing the adaptation time after each environmental transition and increasing the robustness of the phenotype during the constant environment between transitions. We show that surviving lineages evolved bistable, epigenetic switching to adapt quickly in fast fluctuating environments, whereas genetic adaptation was favored in slowly fluctuating environments. For some evolutionary conditions, a strategy emerged where the population adapted genetically between two bistable genotypes.

## INTRODUCTION

Populations often need to adapt to new environments in order to survive. Classic models of genetic adaptation presume the emergence of random genetic mutations, which can generate novel phenotypes that are then selected for by the new environment [1]. Under some circumstances, e.g. small populations or fast fluctuating environments, the spontaneous appearance of a beneficial mutation might be too infrequent and, thus, some organisms may induce genetic variation to accelerate adaptation to a new environment. Examples include adaptive mutation, where genetic variation occurs in response to the environment [2], directed mutation, where non-random and useful mutations are induced [3], and cryptic variation, where non-neutral mutations accumulate and are buffered without phenotypic consequences until the environment changes [4].

Alternatively, many organisms have evolved biochemical networks that directly sense environmental cues and induce adapted phenotypes to the new environment (known as “phenotypic plasticity”). The frequency of environmental change, the accuracy of cues, the penalty of non-adaptation, as well as the cost of producing the sensing machinery will determine the advantage of phenotypic plasticity [5, 6].

A third option is to generate spontaneous phenotypic variation (i.e. not in response to the environment) but restrict this variation to specific phenotypic states. This is possible through the evolution of epigenetic switches, where multiple stable phenotypic states (e.g. level of gene expression) are possible in a genetically identical population and stochastic transitions between these states occur without genetic mutations [7]. This corresponds to the original definition of “epigenetics” by Waddington [8], and has been proposed as one mechanism of bet-hedging to deal with fluctuating environments [9]. Epigenetic switches can emerge through self-reinforcing feedback loops in biochemical or genetic networks, and the epigenetic phenotypes are heritable from mother to daughter cells. Examples of epigenetic switches include heritable gene regulation by self-reinforcing transcription factor activity, DNA methylation, chromatin modification, non-coding RNAs, and prions [10]. Spontaneous transitions between epigenetic states (an “epimutation”) can occur due to biochemical fluctuations and noisy gene expression, where the epimutation rate is usually much faster than genetic mutation [11]. Epimutation generates phenotypic diversity and quick adaptation when the new expression state is favored in the changed environment. As such, epigenetic switches represent an alternative strategy to slower genetic adaptation and costly phenotypic plasticity. Epigenetic switches have been shown to occur in natural populations [12], and to evolve as an adaptation to fluctuating selection during laboratory evolution of microbes [13, 14]. Nevertheless, the specific evolutionary conditions that lead to the selection of epigenetic switches remain to be elucidated.

The advantage of phenotypic plasticity and epigenetic switches in fluctuating environments has been extensively studied using mathematical models and simulations [5, 15–35]. The authors analyzed the effect of different switching rates, either spontaneous or inducible by the environment, on the population growth rate in the context of induction delays, reliability of environmental cues, selection pressures, and environmental dynamics (see Supplementary Table 1). Having phenotypic plasticity was almost always advantageous when the associated cost was low (i.e. small induction delays, environmental cues reliably switch cells to optimal adapted phenotypes, small metabolic burden of biochemical network). In conditions where phenotypic plasticity was disfavored or not available, the authors showed that an epigenetic switch conferred an advantage when the spontaneous epimutation rates matched the environmental fluctuation rate and when selection pressures on fit/unfit phenotypes were symmetric between the two different environments.

These theoretical models did not explicitly include genetic mutation, a competing process that occurs in all organisms. Moreover, many models assumed a non-evolving population and the pre-existence of an epigenetic switch with discrete phenotypes, each optimal in a particular environment. These simplified models often had analytical solutions, however, the absence of population dynamics and genetic adaptation might lead to model-specific idiosyncrasies. For example, many epigenetic switches maximized long-term growth rate by never switching and “ignoring” the environmental transitions [19, 21, 22]. The evolutionary transition from favoring switching vs non-switching was abrupt and exhibited characteristics of a first-order phase transition [22]. However, a non-switching strategy might be destabilized by including competition from a continuous range of genotype-phenotypes accessible through genetic mutation.

A self-activating gene is the simplest genetic circuit that can simultaneously adapt genetically and exhibit epigenetic switching. Recent work by Soyer and colleagues used computer simulation to evolve a self-activating gene in a population of cells surviving in a fluctuating environment [27, 36]. Each generation, they simulated the stochastic gene expression (i.e. phenotype) for each cell with a set of biophysical parameters (i.e. genotype). They modelled the evolutionary process for a population of size *N* using a modified Wright-Fisher model with selection and mutation. Cells with phenotypes that best matched the current environment were preferentially selected to populate the next generation. Each selected cell could mutate its genotype with a fixed probability *u*. The environment fluctuated with frequency *ν* between two discrete states where each environment selected a different gene expression level. The authors concluded from their results that bistability and epigenetic switching were an accidental byproduct of selection for increased nonlinearity.

Here, we used a similar model to simulate the evolution of a self-activating gene over a wide range of selection pressures, mutation rates, mutation step-sizes, population sizes, and environmental fluctuation frequencies. By tracking genotypes and individual lineages that persisted –with or without mutations– across generations, we show that each type of strategy (epigenetic switching or genetic adaptation) was selected under different evolutionary conditions. In fast fluctuating environments, the surviving lineages evolved bistable, epigenetic switching as a strategy, where as genetic adaptation was favored in slowly fluctuating environments. Importantly, we show that epigenetic switching is not an accidental byproduct and that it was selected for in fast fluctuating environments precisely because it adapted more quickly through epimutation. When the mutation step-size (*M*) was small in slowly fluctuating environments, a hybrid adaptation strategy emerged where a bistable population would adapt genetically. We observed a trade-off between minimizing the adaptation time after each environmental transition and increasing the robustness of the phenotype during the constant environment between transitions. The balance between these quantities was modulated by the nonlinearity (Hill coefficient, *n_H_*) in the genetic circuit with smaller values resulting in faster adaptations and larger values resulting in more robust gene expression. We show that each adaptation strategy favors a different aspect of this trade-off: epimutation in epigenetic switching allows faster adaptation, while the adapted genotypes via genetic adaptation are more robust to stochastic noise during the constant environment between transitions. This trade-off helps explain the observed gradual progression between selected adaptation strategies.

## MODELS

The self-activating gene can display a unimodal distribution (monostable) or a bimodal distribution of protein number (bistable), depending on the underlying biophysical parameters (Figure 1 and Figure S1). The spontaneous transitions (i.e. epimutation) between bistable phenotypes are driven by stochastic gene expression. Genetic mutations change the biophysical parameters (i.e. genotype) and will modify both the distribution of protein number (i.e. phenotype, *ρ*(*A*)) and the stochastic transition rate between bistable phenotypes. Two qualitatively different strategies of evolutionary adaptation could emerge from a self-activating gene. The population could evolve from one monostable distribution to another by mutating its biophysical parameters after each environmental change (i.e. genetic adaptation). Alternatively, the population could reside at a bistable solution where each bimodal state is optimal in one of the environments and epimutations from one bistable state to another occur over time without an underlying genetic mutation (i.e. epigenetic switching). In both cases, the phenotypic distribution of the population *P*(*A*) expands each generation due to gene expression noise and mutations, but natural selection keeps it centered on the optimal protein number as determined by the fitness function of the current environment. After an environmental change, the fitness function changes and the tail of the phenotypic distribution with higher fitness will be selected every generation. In the case of genetic adaptation, this selection will gradually shift the population towards the new optimal phenotype through the accumulation of new mutations until the population is well-adapted again. The speed of genetic adaptation depends on the rate of arrival of fitter mutations, as well as the selection pressure. When the population applies epigenetic switching, the phenotypic noise also includes epimutations, which increase the maladapted fraction in the population (epimutational load). However, after the environment and the fitness function change, the “epimutated” individuals rapidly overtake the population, quickly shifting the distribution to the new optimal value. Thus, genetic adaptation and epigenetic switching can directly compete as two strategies of adaptation to a fluctuating environment.

**FIG. 1.**
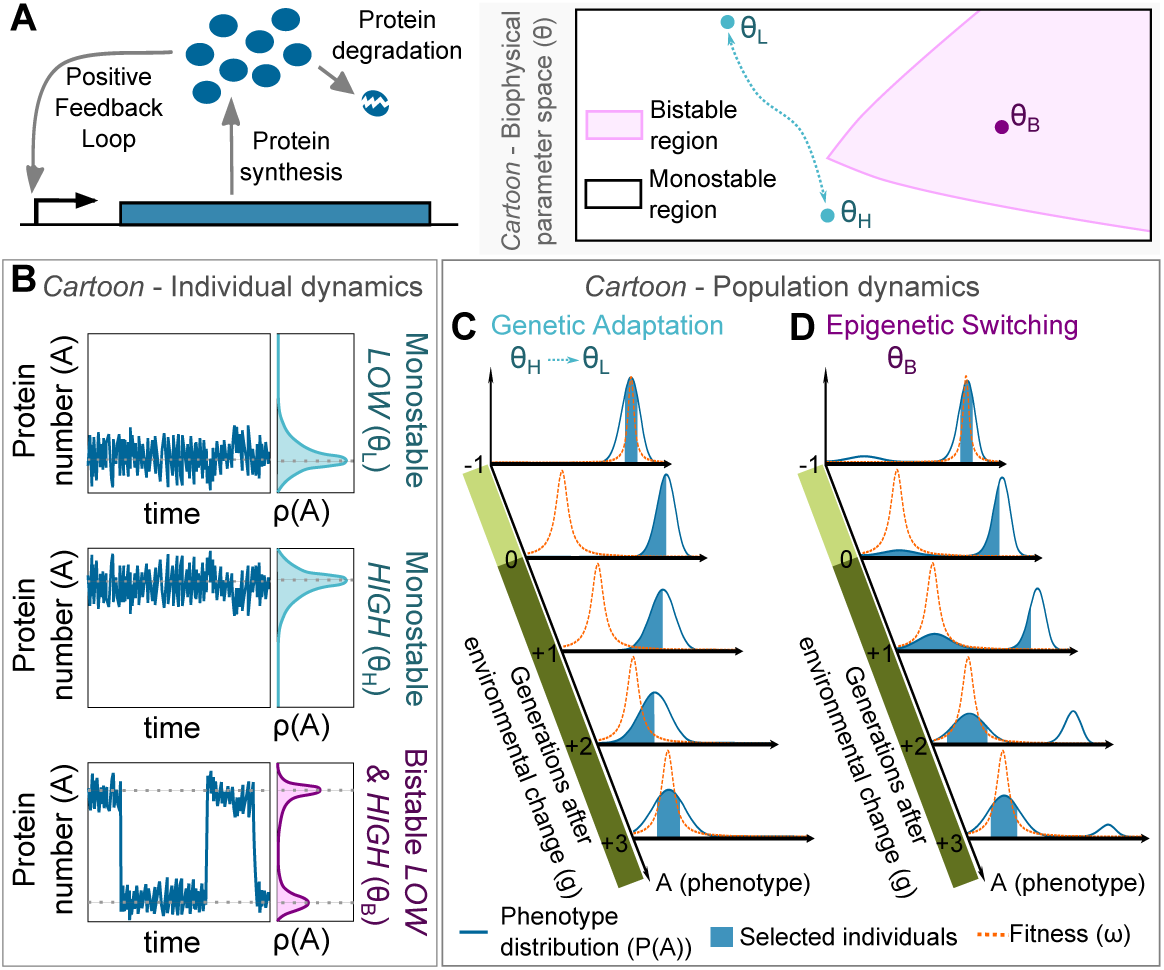
Genetic adaptation versus epigenetic switching of a self-activating gene. (A) Diagram of a self-activating gene that considers two biochemical events: protein synthesis with a rate that increases with number of proteins *A* (i.e., positive feedback loop) and protein degradation. On the right, a cartoon of the biophysical parameter space or genotypes (θ) of this gene circuit with two characteristic regions: monostable (white) and bistable (pink) phenotypes. In the monostable region, a genotype might be optimal in either one environment (e.g. LOW protein numbers, *θ_L_*) or the other (e.g. HIGH protein numbers, *θ_H_*). Genetic mutations are required to change from one solution the other (blue arrow). In the bistable region, a single genotype (e.g. *θ_B_*) can display two different phenotypes with each phenotype potentially optimal in both environments. (B) Cartoon of the protein number (*A*) dynamics in an individual cell with each of the genotypes described in (A). Monostable genotypes (*θ_L_* and *θ_H_*) exhibit a unimodal distribution of protein expression (*ρ*(*A*)), whereas a bistable genotype (*θ_B_*) exhibits a bimodal distribution having spontaneous transitions between phenotypic states over time (i.e. epimutations) triggered by stochastic gene expression. Cartoon of the population dynamics using (C) genetic adaptation or (D) epigenetic switching to adapt after an environmental change. The fitness score function (*ω*; orange dashed line) and the phenotype distribution of the population (*P*(*A*); blue line) are shown for each generation (*g*), and the fraction of the population expected to be selected in the next generation (i.e. individuals with higher fitness scores) are highlighted (blue area). The environment changes from selecting HIGH protein numbers (light green) to select LOW protein numbers (dark green).

### Stochastic dynamics of a self-activating gene

For simplicity, we considered two biochemical events (protein synthesis or degradation) that either increase or decrease the number of proteins *A* by one molecule:

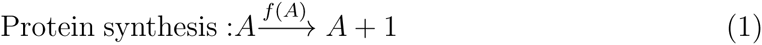

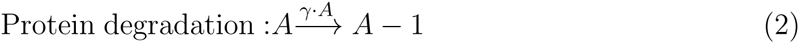

The synthesis rate *f*(*A*), which describes the probability per unit time that protein synthesis occurs and that *A* is increased by one, is a nonlinear function of activator *A*:

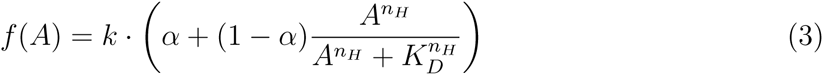

where *k* (proteins/unit time) represent the maximum synthesis rate, α is the basal synthesis rate relative to *k*, *K_D_* (proteins) is related to the protein-DNA dissociation constant, and *n_H_* is the degree of molecular cooperativity (i.e. Hill coefficient). The degradation rate γ · *A*, which describes the probability per unit time that protein degradation occurs and that *A* is decreased by one, is a linear function of activator *A* where γ (1/unit time) is the protein degradation rate constant. With no loss of generality, we defined the unit of time as *τ* = *t·γ*, which allows us to substitute time *t* with a time-dimensionless variable (see *Methods*). We used the Gillespie algorithm, a kinetic Monte Carlo method that explicitly simulates the probabilistic dynamics of a defined set of biochemical events [37], to simulate the stochastic dynamics of our gene circuit (see *Methods*).

### Evolutionary model

For simplicity, we evolved a haploid, asexual population with non-overlapping generations in a fluctuating environment (Figure 2). The underlying biophysical parameters depend on protein stability, protein-protein, protein-RNA, and protein-DNA interactions, which can increase or decrease through genetic mutations. For simplicity, we allowed mutations on the maximum synthesis rate (*k*), Hill coefficient (*n_H_*) and DNA dissociation constant (*K_D_*) during our evolutionary simulations, while keeping basal activity (α = 0.25) and degradation rate (rescaled, γ = 1) fixed. The set of variable parameters {*k, n_H_, K_D_*} are the genotype (*θ*). The environment fluctuated periodically with frequency *ν* between LOW (selects for optimal phenotype *A*^(*L*)^ = 20 proteins) and HIGH (selects for optimal phenotype *A*^(*H*)^ = 80 proteins). The number of generations spent in a constant environment (“epoch”) was equal to 1/*ν* generations. The selective environment switched to the alternative state at the end of an epoch. Starting from an isogenic population, we ran each evolutionary simulation for 10,000 generations. We implemented the following algorithm each generation:

**FIG. 2.**
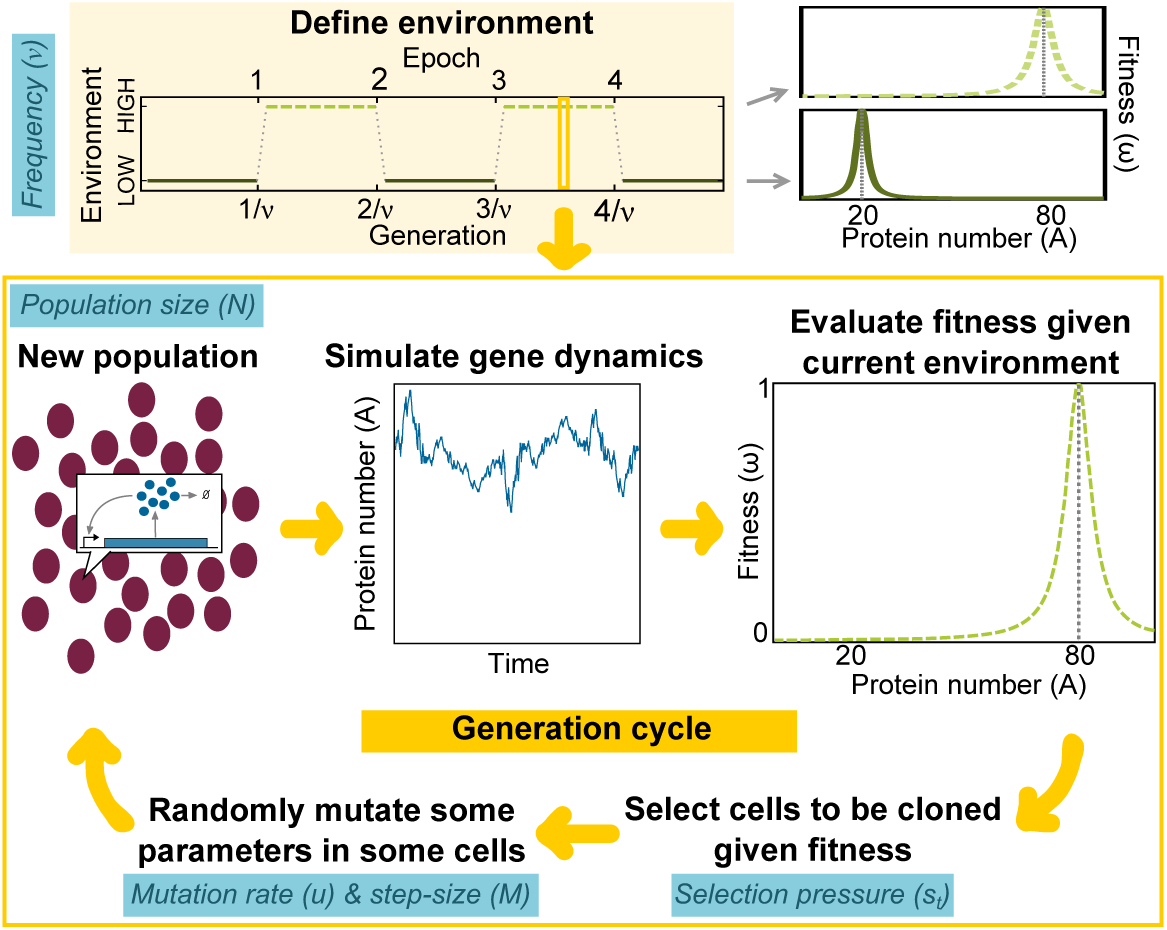
Evolutionary model. The environment fluctuates periodically with frequency *ν*. The total number of generations spent in a constant environment (epoch) has the same length (1/*ν*) and each environment (HIGH or LOW) selects for a different distribution of protein levels (phenotypes). Each generation, we simulated the stochastic protein dynamics of a self-activating gene in each cell across a population of size *N*. At the end of each simulation, the population phenotypes varied because gene expression is stochastic and because cells can have different underlying biochemical parameters (genotypes). The current environment in each generation assigned a fitness (*ω*_*i*_) to each cell (*i*) based on its final protein level. We used Tournament selection (where *s_t_* determines the strength of selection) to determine the next generation of cells according to their fitness. Each cell in the next generation was mutated with probability *u*, where the current set of biophysical parameters were multiplied or divided up to a maximum step-size of *M*.

1. Simulate stochastic gene dynamics of a self-activating gene in each cell given its biophysical parameters (i.e. genotype) and initial protein level inherited from its parent in the previous generation for 4 units of time (the cell “life span”).
2. Evaluate the fitness of each cell *i* based on the protein level at the end of its life span (*A_i_*, phenotype). The individual fitness function (*ω*_*i*_) is:

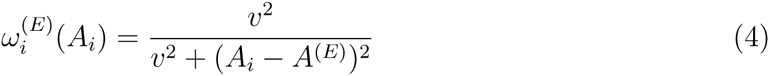

where *E* = {*L, H*} is the current environment, *A* ^(*E*)^ is the optimal phenotype for each environment, and *ν*^2^ = 0.2 · A^(*E*)^ is the width of the Lorentzian function. We define the population fitness (*w*) as the average 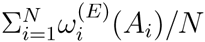 over all cells.
3. Select the next generation using Tournament selection [38, 39], where *s_t_* cells are chosen randomly from the population. The cell with highest fitness within the chosen cohort is cloned into the new population. This “tournament” is repeated *N* times with replacement to create a new population. The tournament size (*s_t_*) modulates the selection pressure, where small *s_t_* is weak selection (e.g. for *s_t_* = 1, there is no selection pressure and only genetic drift because any randomly selected individual is the tournament winner). Increasing *s_t_* leads to stronger selection and a faster selective sweep of fitter cells (e.g. for *s_t_* = *N*, only the fittest individual in the entire population will be cloned into the next generation).
4. Allow random mutations with a fixed probability (*u*) in each cloned cell. If a mutation occurs, the value of each parameter in the cell genotype is updated as follows:

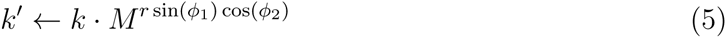

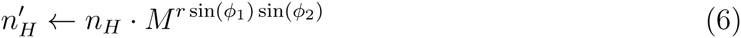

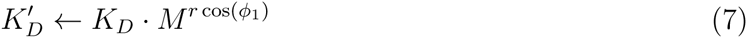

where *M* is the maximum fold change (mutation step-size), *r* ∼ *U* (0, 1) is a uniformly distributed random value between 0 and 1, *ϕ*_1,2_ ∼ *U* (0, 2*π*) are uniformly distributed random values in radian units. Unlike additive mutation, multiplicative mutation better reflects how mutations affect thermodynamics of protein stability, protein-DNA and protein-protein interactions [40, 41]. All mutated parameters were constrained to lie within a physiological range that is typical for a bacterial transcription factor (see *Methods*).
5. If the evolutionary simulation is at the end of an epoch, then change to other environment; otherwise keep the same environment. Return to Step 1 to simulate next generation.

## RESULTS

The evolutionary model was simulated over a range of population size (*N* = [100, 10000]), selection pressure (*s_t_* = [3, 250]), environmental fluctuation frequency (*ν* = [0.01, 0.1]), mutation rate (*u* = [0.01, 0.1]), and mutation step-size (*M* = [1.1, 5]). We restricted these parameters to regions where our simulations were feasible and where epigenetic switching and genetic adaptation were competitive with one another. For example, we only considered *uN* ≥ 1; otherwise, mutations were too infrequent for genetic adaptation to compete with epigenetic switching. We verified that our results presented below were robust to alternative assumptions, such as different models of selection, non-periodic environmental fluctuations, a Moran model of reproduction, and alternative mutation schemes (see *Methods*).

To control for the possibility that bistability could be selected for reasons other than epigenetic switching [27], we also ran a parallel deterministic simulation (CONTROL) of the expression of the auto-activating gene in Step 1 (see *Methods*). Given that there is no biochemical noise in our CONTROL simulations, cells in the bistable region display hysteresis and stay in the stable state closest to the inherited parental state and never stochastically switch to the other state (i.e. no epigenetic switching).

### Epigenetic adaptation to fluctuating environments

All populations initially started with a monostable genotype ϕ_0_ that was adapted to one environment but not the other. These populations all evolved to a higher fitness solution and strategy that was best suited for the underlying evolutionary parameters. For example, in a fast-fluctuating environment (*ν* = 0.1) with small mutation step size (*M* = 1.1), the population evolved to a bistable genotype *θ_B_* that used epigenetic switching. Initially, the population had a slightly higher geometric mean fitness per cycle 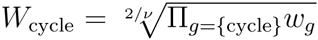 because it accrued benefits by being adapted to the HIGH environment despite being mal-adapted to the LOW environment (Figure 3). We plot geometric fitness per cycle rather than fitness per generation (*w*) because it better reflects the long-term growth of fitter phenotypes [42]. During the LOW epoch that follows the HIGH epoch, the population shifted towards higher *w*^(*L*)^ values. The epoch was too short and mutation too weak for the population to perfectly adapt to the new environment before it changed again. The evolutionary dynamics in early epochs were dominated by noisy genetic adaptation of a population maladapted to at least one of the environments, even if this implied decreasing *W*_cycle_ (Figure S2). The “no-response” behavior, i.e. being adapted to one environment and “ignoring” the alternative state, is not a stable solution for this system. Similar to previous work, this illustrates the importance of considering the full population dynamics in the adaptation process and not only the long-term average fitness [43].

**FIG. 3.**
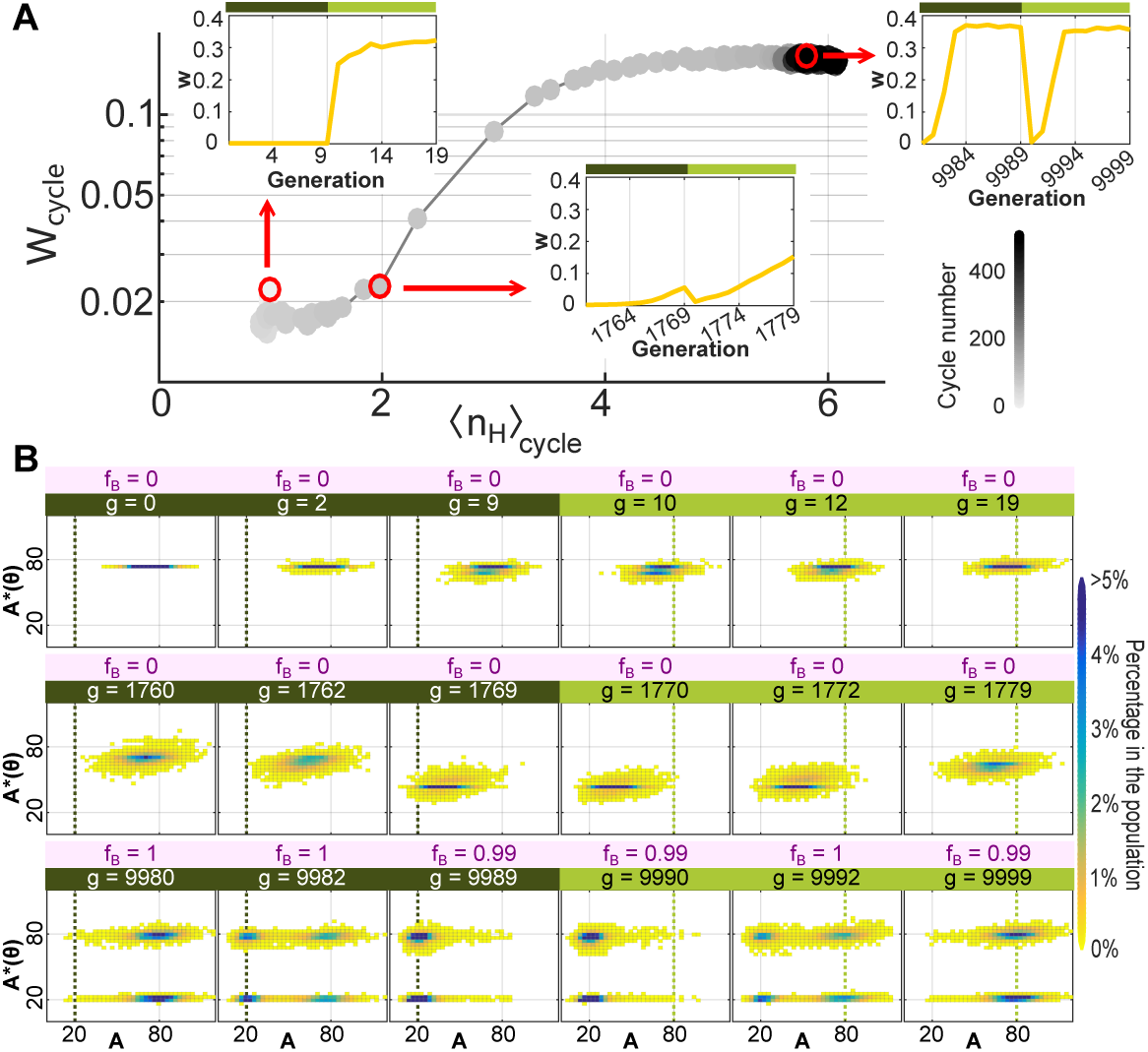
Evolutionary trajectory of a population adapting to a fast fluctuating environment. The initial population started from a non-optimal genotype (θ_0_) where *k* = 80, *n_H_* = 1, *K_D_* = 10 with evolutionary parameters *N* = 10000, *ν* = 0.1, *s_t_* = 6, *u* = 0.03, and *M* = 1.1. (A) The geometric mean of the population fitness per environmental cycle (*W*_cycle_) versus the average Hill coefficient per cycle (〈*n_H_*〉_cycle_). Each cycle spans a LOW (dark green) and HIGH (light green) epoch and there are 500 environmental cycles (increasing from gray to black) over 10,000 generations for this simulation. The initial genotype was well adapted only to the HIGH environment (see population fitness *w* per generation in inset), but the forces of mutation-selection in the alternative environment gradually shifted the population to an intermediate genotype with higher 〈*n_H_*〉_cycle_ that lay between the two optimal phenotypes for each environment (see inset). The population then evolved to the bistable parameter space with higher 〈*n_H_*〉_cycle_ and *W*_cycle_. This final genotype had high population fitness (*w*) in both environments and rapidly adapted after each environmental transition through epigenetic switching (see inset). (B) Population dynamics for initial, intermediate, and final cycles highlighted in (A) are shown. At each generation, we plot the joint distribution of the actual phenotype (*A*) from stochastic simulation and the expected steady-state phenotype(s) *A*^*^(θ) from the deterministic solution given the individual’s genotype. The optimal phenotype given the environment is shown as dotted line, and the fraction of bistable individuals in the population (*f_B_*) is listed in the top.

The sub-optimal population eventually increased the nonlinearity in gene expression 〈*n_H_*〉_cycle_ before transitioning from a monostable to a bistable genotype at *∼* 1800 generations (Figure 3A). The bistable genotype was a global optimum in *W* cycle and adaptation occurred in a few generations after each environmental change due to epigenetic switching. We verified that this bistable genotype was globally optimal by re-running evolutionary simulations for different initial genotypes (*θ*_0_) and for more generations (data not shown). The epigenetic switching can be visualized by plotting the actual phenotype *A* from stochastic simulation versus expected deterministic phenotype *A*^*^(*θ*) of the evolving population over an environmental cycle (Figure 3B). Bistable cells have two possible deterministic phenotypes *A*^*^(*θ*) as shown on the y-axis. Our data show that the final population adapted to the fluctuating environment by finding a genotype (*θ_B_*) in the biophysical parameter space where the transition between bistable solutions was efficient.

### Genetic adaptation to fluctuating environments

Monostable populations can adapt genetically through mutation after a change in environment (i.e. random walk in parameter space *θ* to a new monostable phenotype; see Figure 1). If mutations are frequent and mutation step-size (*M*) is large, then genetic adaptation can occur within several generations and outcompete bistable epigenetic switching. Monostable populations have potentially lower gene expression noise cost than bistable populations (which have an increased fraction of maladapted phenotypes arising from epimutations). Thus, longer epochs in a constant environment (smaller ν) should also favor genetic adaptation over epigenetic switching.

To explore the transition from epigenetic switching to genetic adaptation, we simulated the dynamics of evolutionary adaptation for longer epochs (smaller *ν*) and bigger mutation step-size (larger *M*). As expected, the fraction of monostable solutions increased in these conditions. However, the fraction of bistable individuals in the population (*f_B_*) never became zero and even showed regular temporal and environmental patterns (Figure 4 and Figure S3). The monostable sub-population shifted their steady state values towards the given optimal phenotype each time the environment changed; while, the bistable sub-populations kept their expected steady-state phenotypes *A*^*^(*θ*) centered on the two optimal values, regardless of the current environment. This co-existence is intriguing because these sub-populations apply different adaptation strategies. If one strategy is more fit than the other, then we would expect one to fix and the other to go extinct. However, longer simulations displayed the same pattern and a larger population size stabilized the observed bistable fraction *f_B_* (Figure S3). In cases where a subpopulation would go extinct, it would re-appear and re-establish itself in subsequent generations (Figure S3). This suggests that “seeding” of one sub-population from another through genetic mutation plays a role in the evolutionary stability of co-existing sub-population fractions.

**FIG. 4.**
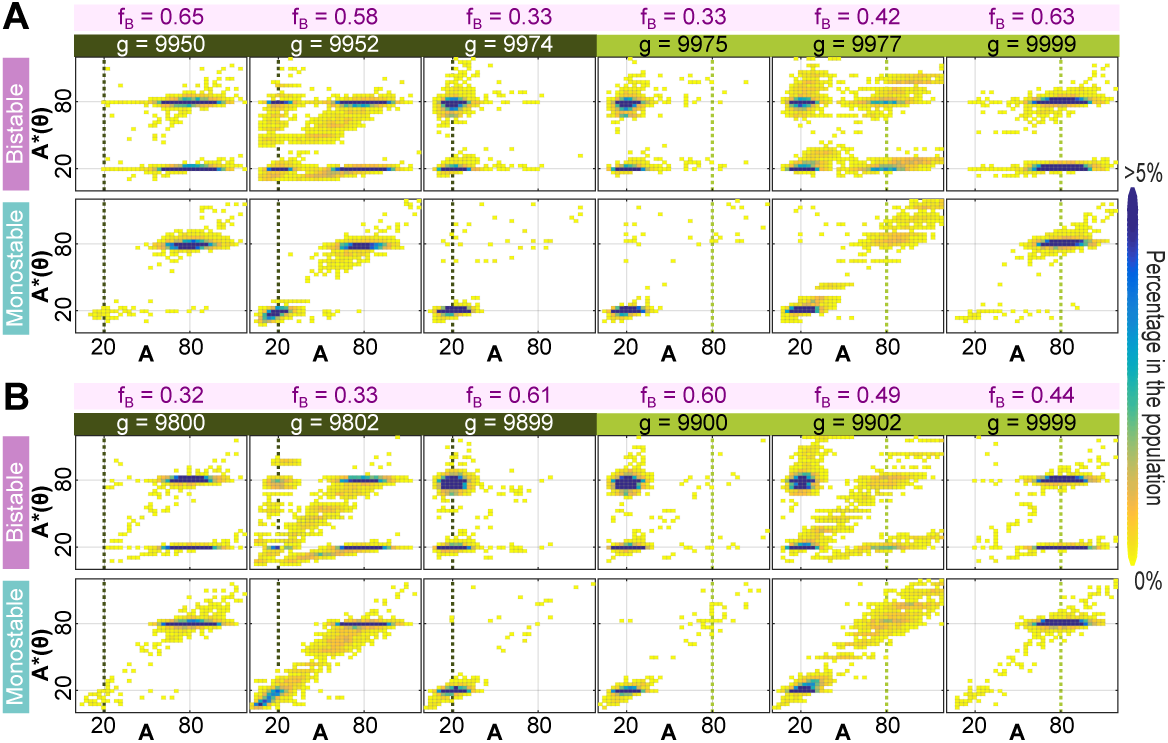
Coexistence of bistable and monostable sub-populations in the final adapted population under multiple evolutionary conditions. Population structure and evolutionary dynamics of adaptation to two environmental states (LOW, dark green; HIGH, light green) at 10,000 generations. Each simulation started from the same genotype and evolutionary parameters as Fig. 3 except (A) *ν* = 0.04, *M* = 2.1 (25-generation epoch, moderate mutation step-size) and (B) *ν* = 0.01, *M* = 5.0 (100-generation epoch, large mutation step-size). The final adapted population contained both monostable and bistable sub-populations with bistable fraction *f_B_* (shown in pink). For each sub-population (bistable, top; monostable, bottom), we plot the joint distribution of the actual phenotype (*A*) and deterministic phenotype *A*^*^(*θ*). As a reference, the optimal phenotypes for each generation are shown as a dotted line.

### Lineage analysis demonstrates that one adaptation strategy is usually dominant each cycle

To better understand the adaptation process and evolutionary forces that generate coexistence, it was informative to analyze the genealogy (i.e. lineages) of the current population (see *Methods*). We tracked the evolutionary history of the population to elucidate those lineages that persisted with or without mutations over one full environmental cycle (i.e. LOW epoch + HIGH epoch; Figure S4). If a particular adaptation strategy is successful, then we expect those lineages using that strategy to have a higher fitness and to persist over multiple environmental cycles. More than one lineage can persist over a cycle, but fewer than expected from coalescent theory because our population is evolving under selection and faces a bottleneck at each environmental transition. The weight of each persisting lineage is proportional to the number of progeny at the end of the cycle.

An epigenetic switch can adapt with no mutation; thus, lineages with a bistable genotype and no mutations during a cycle were classified as having an epigenetic switching (ES) strategy (Figure 5A). On the other hand, those lineages that had at least one monostable genotype and accumulated mutations during a cycle were classified as having a genetic adaptation (GA) strategy. Lineages with only bistable genotypes that accumulated mutations during a cycle were classified as having a hybrid bistable adaptation (BA) strategy. Although some of these mutations can be neutral, most of them modulated the DNA dissociation constant (*K_D_*) and directly affected the rate of epigenetic switching (“epimutation”); see Figure 5A. Thus, the mutation can be adaptive in the hybrid BA strategy, although the circuit remained bistable and ultimately adapted through epigenetic switching.

**FIG. 5.**
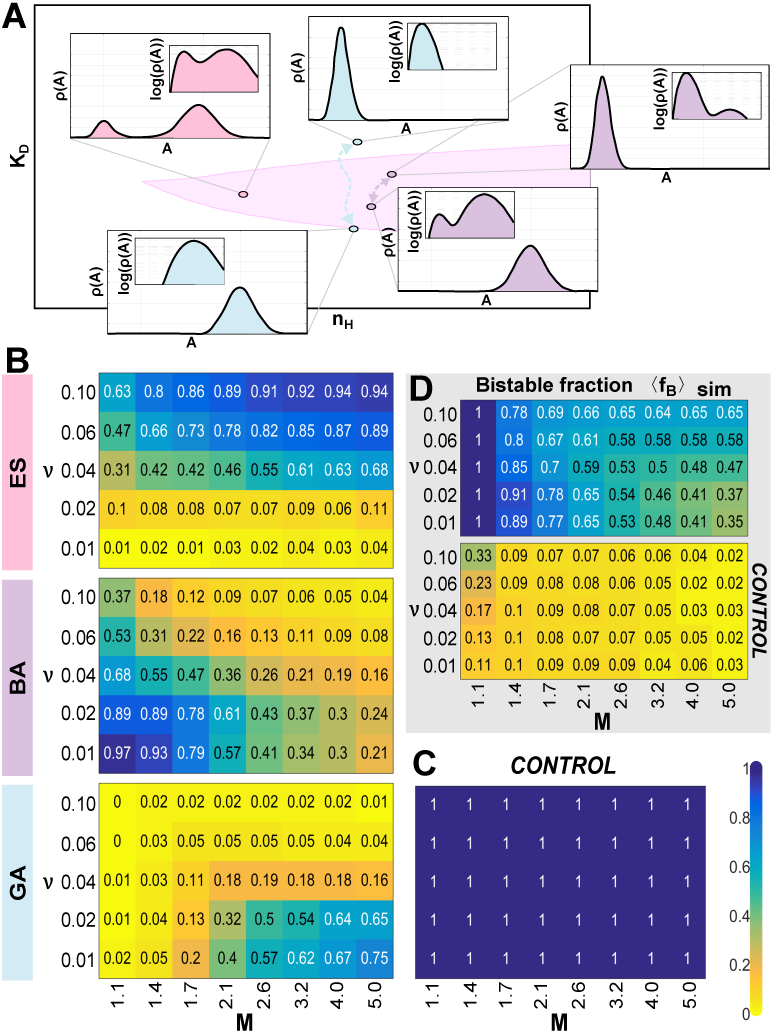
Each adaptation strategy is favored under different evolutionary conditions and the transition between selected strategies is gradual. (A) Illustration of epigenetic switching (ES), bistable adaptation (BA), and genetic adaptation (GA) strategies and underlying genotypes with fixed synthesis rate (*k* = 80). The bistable region of genotype space is highlighted in pink. The phenotype distribution *ρ*(*A*) for each genotype (*θ*) is shown in the inset, both in linear and logarithmic scale. BA is bistable, as seen in logarithmic scale, but appears effectively monostable in linear scale. This arises because *K_D_* evolves each epoch to favor one mode over the other by decreasing the relative rates of epigenetic switching between the largest and smallest mode. (B) Each colormap shows the fraction of parental lineages using specific adaptation strategy (ES, BA, or GA) averaged over all cycles and ten independent replica simulations for the corresponding mutation step-size (*M*) and environmental fluctuation frequency (*ν*). Each simulation ran for 10,000 generations with evolutionary parameters *N* = 10000, *s_t_* = 6, *u* = 0.03 and *k* = 80, *n_H_* = 6 and *K_D_* = 45 as the initial genotype (*θ*_1_). This initial genotype sped up evolutionary simulations by being closer to final selected genotypes in all simulations. (C) Results of the CONTROL simulations where gene expression dynamics are deterministic and no stochastic epigenetic switching can occur. All lineages exhibited GA and neither bistable strategy (ES or BA) was ever selected. (D) The corresponding bistable fraction (〈*f_B_*〉_sim_) averaged over all cycles and ten independent replica simulations for the stochastic simulations (top) and deterministic CONTROL (bottom).

Lineage analysis demonstrated that distinct strategies were favored in different conditions (Figure 5B). If the environment fluctuated frequently (i.e. high *ν* values), the dominant adaptation strategy was ES. In slowly fluctuating environments (i.e. low *ν* values), GA was used if the mutation step-size (*M*) was large enough; otherwise, BA was the dominant adaptation strategy. Interestingly, we observed a mixture of strategies across lineages (no strategy was 100%), and the transition between preferred adaptation strategies as a function of *ν* and *M* was gradual. These trends persisted over a range of different evolutionary parameters, although the location of the boundaries would shift. For example, increasing the selection pressure (*s_t_*) or the mutation rate (*u*) shifted boundaries to favor GA, whereas increasing population size (*N*) favored ES (Figure S5).

Our CONTROL simulations with deterministic dynamics (where no stochastic epigenetic switching can occur even if the system is bistable) showed that none of the parental lineages exhibit ES or BA; see Figure 5C. The preference for GA in simulations with deterministic dynamics demonstrates that ES and BA are selected for by stochastic simulations precisely because of their ability to spontaneously switch phenotypes (epimutation).

We also investigated the average fraction of bistable genotypes (〈*f_B_*〉_sim_) as a function of evolutionary parameters for stochastic simulation and deterministic CONTROL (Figure 5D). In broad agreement with Figure 4, 〈*f_B_*〉_sim_ never decreased to zero even when GA was the preferred lineage strategy (e.g. low *ν*, high *M*). These results arise from the increased seeding of genetic mutants (which is facilitated by higher *M*) from the monostable to bistable subpopulation. Conversely, we expect more cases of neutral or near neutral mutations in the bistable region for smaller mutational step-sizes (*M*). It is exactly in this regime where the fraction of ES parental lineages decreased whereas BA increased. However, most lineages displaying BA in this regime also persisted into the next cycle without accumulating any mutations and, thus, automatically switched to ES (see Figure S6 for rates of lineage switching from one strategy to another each cycle).

### Fitness reflects a trade-off between adaptation time and phenotypic robustness

Our simulations showed that different portfolios of strategies are favored for different mutation step sizes (*M*) and environmental transition frequencies (*ν*). Bistable strategies (ES or BA) require genotypes with large nonlinearity (high *n_H_*), whereas GA genotypes are presumably less constrained. However, all evolutionary simulations (including GA) evolved to average genotypes with high nonlinearity 〈*n_H_*〉_sim_ (Figure 6A). This trend persisted in deterministic CONTROL simulations where ES and BA cannot occur (data not shown). The evolution to genotypes with high nonlinearity is consistent with previous work [27]; see *Discussion* for our explanation why all simulations evolved to genotypes with high nonlinearity. Strikingly, 〈*n_H_*〉_sim_ was mostly determined by the environmental fluctuation frequency and it increased as *ν* decreased (i.e. longer epochs; Figure 6A). This same trend was observed when we measured the geometric mean fitness averaged over the whole simulation 〈*W*_cycle_〉_sim_ (Figure 6B). Populations that evolved in slowly fluctuating environments had higher average fitness and higher nonlinearity.

**FIG. 6.**
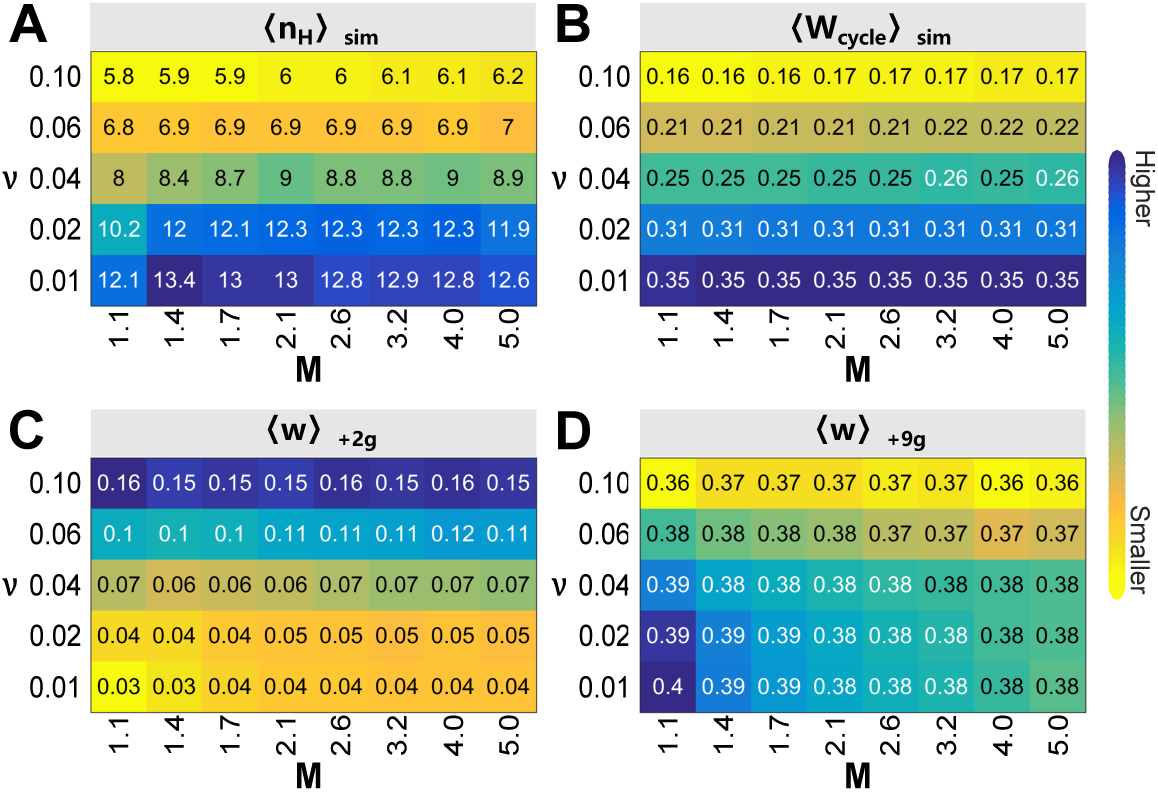
Population fitness depends on environmental fluctuation frequency and reflects a trade-off between adaptation time and phenotypic robustness. (A) Hill coeffi-cient 〈*n_H_*〉_sim_ averaged over ten independent replica simulations for mutation step-size (*M*) and environmental fluctuation frequency (*ν*). (B) Average geometric mean fitness per cycle 〈*W*_cycle_〉_sim_ for the same simulations. (C) Average population fitness at the second generation *〈*w*〉*_+2*g*_ and (D) ninth generation *〈*w*〉*_+9*g*_ after an environmental transition. Evolutionary strategies with a faster adaptation time had larger fitness after the transition (C) whereas those with robust phenotypes tended to have a larger fitness once they adapted to the new environment (D). All simulations ran for 10,000 generations with identical evolutionary parameters and initial genotypes as in Figure 5.

The geometric mean fitness per cycle, which includes the lower fitness of the transient, maladapted population immediately after a transition in addition to the higher fitness of the adapted population, was surprisingly insensitive to the mutation step-size (*M*). One might expect that a larger mutation step size increases the mutational load and, thus, decreases the average fitness of the final adapted population. This discrepancy suggested to us that the population fitness cost of new mutations (mutational load) was minimal for our mutation rate (*u*) when compared to other costs (e.g. gene expression noise and/or epimutational load). In agreement with this hypothesis, increasing the mutation rate (*u*) generated *M*-dependence in fitness, whereas decreasing the mutation rate had little effect (Figure S7). Varying the selection pressure (*s_t_*) only affected the actual 〈*W*_cycle_〉_sim_ values observed, but not the trend (Figure S8). Similarly, varying the population size (*N*) had no significant effect on the population fitness behavior, except that results became noisier as *N* decreased (Figure S9).

The trend in average geometric mean fitness 〈*W*_cycle_〉_sim_ as a function of can be understood by considering the timescales of epochs. The population fitness is always low and maladapted when the environment first changes. Selection favors those genotypes which produce phenotypes that better match the selection pressure in the new environment. In this first phase (the “adaptation phase”), the fitness increases as the population adapts through genetic adaptation (GA and BA, mutation) and/or epigenetic switching (BA or ES, epimutation). The average population fitness two generations after each environmental change 〈*w*〉_+2*g*_ is *M*-independent and highest for the ES genotypes that emerge for large ν (Figure 5B and Figure 6C). The rate of epimutation is faster than genetic mutation and, thus, ES (the dominant strategy) adapts more quickly and has a higher transient fitness. However, all populations reach a higher fitness and become adapted to the new environment. In this second phase (the “constant phase”), purifying selection maintains the optimal phenotype against perturbations from gene expression noise, mutations, and/or epimutation. The average population fitness nine generations after each environmental change 〈*w*〉_+9*g*_ is mostly *M*-independent and highest for small *ν* (Figure 6D). The population spends proportionally more time in the constant phase as *ν* decreases, which favors those strategies with robust phenotypes (i.e. minimize frequent, maladapted phenotypes that arise from epigenetic switching). Thus, 〈*W*_cycle_〉_sim_ increases and GA or BA genotypes emerge as decreases because the population spends more time in the “constant phase” and epigenetic switching does not occur or is effectively zero (see *ρ*(*A*) in Figure 5A).

These results suggest a trade-off in the evolutionary process between minimizing the adaptation time during the adaptation phase and increasing the robustness of the phenotype during the constant phase (Figure 6). This trend correlates with the selected Hill coefficient value in the population 〈*n_H_*〉_sim_, which suggests that natural selection tunes this trade-off via this biophysical parameter (Figure S10); see *Discussion*. The trade-off between two fitness costs also explains the observed gradual transition between selected adaptation strategies (Figure 5B). These fitness costs are continuous and can have very similar values, such that genetic drift will dominate during the selection process. As expected, the simulations in regimes with co-dominant strategies showed a high temporal variation in the fraction of adaptation strategies each evolutionary cycle (Figure S11).

## DISCUSSION

### Epigenetic switches are selected by fast fluctuating environments

Previous theoretical work established that optimal long-term growth occurs when the phenotype switching rate matches the environmental switching rate [15–18]. The phenotype could switch either due to genetic adaptation with a rate that depends on the mutation rate and mutation step-size, or due to epigenetic switching (epimutation) with a rate determined by the underlying molecular system. In the natural world, epimutation rates are often faster than genetic mutation rates [11], which suggests that fast fluctuating environments might select for epigenetic switching (ES) over genetic adaptation (GA). Previous models did not integrate and evaluate these two competing processes in a population of cells evolving in a fluctuating environment.

To this end, Soyer and colleagues used computer simulations to evolve a population of self-reinforcing gene circuits (which can exhibit epigenetic switching and genetic adaptation) in a fluctuating environment [27, 36]. Their work demonstrated the emergence of ES (bistable genotypes) in fast fluctuating environments. However, the authors proposed that ES was accidental and that bistability emerged as a byproduct of selection for increased nonlinearity and higher evolvability (i.e. large changes in phenotype with small changes in genotype).

Our simulations confirm that ES emerges in fast fluctuating environments across many different conditions (Figure 5). In agreement with Soyer and colleagues, our populations also evolved to genotypes with high nonlinearity (Figure 6A). However, we disagree that ES is an accidental byproduct of selection for increased nonlinearity and evolvability. Bistable genotypes were only favored in the presence of gene expression noise when cells can stochastically switch phenotypes (i.e. non-zero epimutation rate). In CONTROL simulations (i.e. deterministic dynamics with no noise and, hence, no epimutation), we only observed GA (Figure 5C). We conclude that ES is selected over GA in fast fluctuating environments precisely because of the benefits of epimutation.

### Larger nonlinearity increases fitness and regulates the trade-off between adaptation time and phenotypic robustness

Starting from an arbitrary initial condition and different evolutionary conditions, all populations evolved to a similar region of genotype space with higher fitness (data not shown). These final genotypes often had *k* ∼ 80, 20 < *K_D_* < 80, and large *n_H_*. To help understand the forces that select for large nonlinearity, we considered a simple model of an infinite, clonal population with steady-state protein distribution *ρ*(*A*) determined by *n_H_*, *K_D_* and fixed *k* = 80. We calculated the expected fitness in LOW and HIGH environment as a function of *n_H_*, *K_D_* (Figure S10). Our simple model demonstrates that the fitness of genotypes with steady state phenotypes *A*^*^(*θ*) close to the optimal values always increases with *n_H_* regardless of *K_D_*, the specific environmental state (LOW or HIGH), or whether the genotype is bistable.

The increased fitness arises from better buffering of gene expression (output) against intrinsic biochemical noise in the protein levels (input). As the nonlinearity increases, the gene expression rate *f*(*A*) both above and below *K_D_* becomes more zero-order and less sensitive to fluctuations in protein levels (*A*). For the same reason, mutations in *K_D_* are less likely to affect steady-state protein levels and will have a smaller effect on fitness. Thus, increasing nonlinearity leads to higher fitness because of increased robustness of gene expression to both biochemical noise and some *de novo* mutations. This connection between nonlinearity, robustness to noisy gene expression and robustness to mutations may be an interesting example of plastogenetic congruence [36, 44, 45].

The average population fitness was mostly determined by the environmental fluctuation frequency (*ν*), where fitness decreased as the epoch length decreased (Figure 6). This agrees with previous theoretical work, which showed that the evolutionary dynamics are governed by environmental dynamics [15, 30]. In our simulations, the correlation between fitness and epoch length arises from the balance between two competing fitness costs: reducing the time required to adapt every time the environment changes (adaptation phase) and increasing the phenotypic robustness when the environment is fixed (constant phase). The ES strategy is selected at high *ν* (when the population spends proportionally more time in the adaptation phase) because it has a faster adaptation time (Figure 6C) at the cost of lower phenotypic robustness (Figure 6D). Our work suggests that the trade-off between adaptation time and phenotypic robustness is mostly modulated by the nonlinearity. A smaller *n_H_* increases the stochastic transition rates of ES at high *ν* and a larger *n_H_* increases the phenotypic robustness of GA at low *ν*.

### Lineages reveal hidden selection forces

All evolved bistable and monostable genotypes were relatively close to each other (Figure 5A). Our simulations had a relatively high mutation rate (*u*) such that bistable genotypes could mutate to monostable genotypes, and vice versa. The elevated rate of seeding between these subpopulations made it challenging to distinguish whether ES (bistable) was being selected for. It has been shown previously that individual history or genealogy can efficiently reveal hidden selection forces. For example, Kussell and colleagues [33, 46] demonstrated that selective pressures on a population, such as those imposed by a fluctuating environment, can be efficiently quantified by measurements on the surviving lineages. More recently, Cerulus et al. [47] used life-history traits of cellular growth to show that high single-cell variance in growth rate can be beneficial for the population, and that this benefit depends on the epigenetic inheritance of the growth rate between mother and daughter cells.

To this end, we analyzed the strategy of lineages across multiple cycles during our simulations. Lineage analysis demonstrated that apparent coexistence of bistable and monostable subpopulations was a transient phenomenon, and one type of strategy was typically dominant across lineages (Figure 5). Our analysis suggests that population snapshots (e.g. bimodal versus unimodal distribution of phenotypes) can miss the contribution of epigenetic switching. Future experimental studies on the evolution of epigenetic switches might consider analyzing lineages using time-lapse microscopy, as done by Balaban et al [12].

### Model limitations and future directions

We verified that our observations were robust to many alternative model assumptions (Figure S12). Nevertheless, our simple model of a haploid, asexual population [48] omits some features of the evolutionary process. For example, variable population size, diploid genetics, sexual reproduction and linkage disequilibrium could all affect the evolutionary dynamics and selection of epigenetic switches in a fluctuating environment. Our model also fixed the mutation rate (*u*) and step-size (*M*), which imposes a mutational load when the population is adapted to a constant environment. Future simulations could allow natural selection to mutate and tune these parameters, which might favor genetic adaptation over epigenetic switching [49]. Our model also did not consider the case where mutations (e.g. adaptive mutation) or biophysical parameters (e.g. phenotypic plasticity) directly respond to changes in the environment. Last, we assumed that mutations continuously increase or decrease the biochemical parameters. This overlooked an important class of mutations, such as indels (i.e. rapid loss or abrupt change of function), gene duplication, and gene recruitment, which could abruptly change the topology of the gene network.

We considered the simplest genetic circuit that can exhibit epigenetic switching. However, alternative gene regulatory networks could generate different dynamics and phenotypes that are even better adapted to the fluctuating environment. For example, adding a negative feedback loop could reduce gene expression noise [50, 51] or generate oscillations [52, 53]. An oscillatory gene circuit (e.g. circadian clock) might anticipate and respond to an environment that fluctuates regularly (e.g. day/night). Future research will explore more complicated gene regulatory circuits to understand the specific environmental dynamics and evolutionary conditions that favor oscillation versus epigenetic switching in the context of genetic adaptation. This should be of broad relevance to evolutionary biologists and systems biologists.

## METHODS

### Biophysical parameters

The maximum synthesis rate *k* and the degradation rate *γ* depend on time. With no loss of generality, we reduced the number of free parameters in our model by substituting time *t* with a time-dimensionless variable *τ* = *t* · *γ*. Many proteins in bacteria are not actively degraded and are diluted through cell growth and division. Thus, *τ* and *k* are in units of cell cycle time. All mutated parameters were constrained to lie within a physiological range (10^−2^ ≤ *k* ≤ 10^3^, 10^−2^ ≤ *n_H_* ≤ 16, 10^−2^ ≤ *K_D_* ≤ 120) for a bacterium such as *E. coli*. The number of molecules for a transcription factor ranges between 0 – 10^3^ proteins per bacterium or concentration range 0 – 10^3^ nM for a bacterial volume of 1 fL [54, 55]. The DNA dissociation constant (*K_D_*) has a similar range to the underlying transcription factors [55, 56].

### Gillespie algorithm

The propensity or probability rate (*r_j_*) of chemical reaction *j* occurring during the next interval *dt* is related to the rates of mass-action chemical kinetics in a constant chemical reactor volume *V*. In our simple biochemical network, the propensity of protein synthesis is *f* (*A*) and the propensity of protein degradation is *γ · A*. Each step, given the current number of chemical species, Gillespie’s direct algorithm first calculates the propensities *r_j_* and then calculates when the next reaction occurs and which one occurred. The waiting time of the next reaction is drawn from an exponential distribution with parameter Σ_*j*_*r*_*j*_, where the cumulative distribution function of any reaction occurring before time *τ* is *F*(*τ*) = 1 − *e*^−(Σ_*j*_*r*_*j*_)·τ^. Given that a reaction has occurred at time *τ*, the probability that the event is reaction *i* is equal to *r_i_*/(Σ_*j*_*r*_*j*_). Thus, for each iteration, two random numbers are required to determine *τ* and *i* as drawn from the probability distributions. The random numbers are generated using the “Minimal” random number generator of Park and Miller with Bays-Durham shuffle and added safeguards [57].

### Deterministic CONTROL simulations

The dynamic of the mean concentration (*A*) obeys a first-order ordinary differential equation:

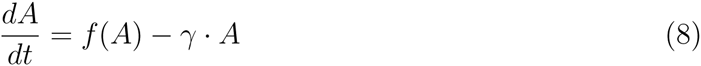

The cell life span was assumed to be long enough for steady state (where Eq. 8 equals 0) to be reached before selection. We used numerical methods to calculate the steady state solution (phenotype) for any genotype and initial protein level inherited from the parent. All Gillespie, deterministic, and evolution simulations were implemented in C++ (*Supplementary code*), and all the analyses and figures were done using MATLAB.

### Lineage analysis

An unfit mutant could arise at the end of a cycle or a fit genotype might go extinct due to genetic drift and gene expression noise. To obtain insights on the evolutionary stability of different strategies, we focused our analysis on lineages that have persisted –with or without mutations– through at least one full environmental cycle (i.e. LOW epoch + HIGH epoch). We maintained genealogical records and traced the lineage and ancestral genotype of all cells over 2 cycles (Figures S4). If *any* genotype between the 1-cycle or and 2-cycle ancestors was monostable, then we classified the evolutionary strategy of that lineage as genetic adaptation (GA). If all genotypes between the 1-cycle and 2-cycle ancestors were bistable, then we classified the evolutionary strategy as either epigenetic switching (ES) or bistable adaptation (BA). The lineage was BA only if the 1-cycle bistable ancestor had accumulated at least one mutation since the 2-cycle bistable ancestor. At the end of each cycle, we calculated the fraction of surviving cells whose ancestors had one of 3 strategies (GA, ES, BA) and averaged over all cycles (Figure 5). Each individual ancestral lineage was counted regardless of whether it was shared with other individuals or not. To determine the transition rate between evolutionary strategies, we analyzed lineages back to the 3-cycle ancestor. We calculated the statistics of transitions between adaptation strategies by comparing strategies in 1-cycle and 2-cycle ancestors (current adaptation strategy) versus 2-cycle and 3-cycle ancestors (previous adaptation strategy) and averaged over all cycles (Figure S6).

### Alternative model assumptions

We tested the robustness of our results to alternative choices and assumptions in the evolutionary model.

#### Environmental fluctuations

Our fluctuations were regular and periodic with frequency *ν*. We tested whether stochastic fluctuations with frequency *ν* produced different results, even though previous work demonstrated little difference between the two types of fluctuations [5, 16, 18, 19]. Our simulations confirmed that periodic and stochastic environmental fluctuations generate the same qualitative trends (Figure S12).

#### Selection algorithms

We used Tournament selection to select the next generation of cells based on the fitness of the individuals in the current generation. Other common selection schemes are Truncation, Proportional, and Weighted selection [58]. These selection schemes produced similar results to Tournament selection (Figure S12). We also obtained similar results with a Moran model (Figure S12), where the birth and death events are treated as continuous, stochastic events instead of non-overlapping generations (as in our modified Wright-Fisher model). The Moran simulations were computationally expensive, which is why we settled on Tournament selection with a Wright-Fisher model.

#### Fitness of phenotypes

Our simulations evaluated the protein number (phenotype) at the end of Gillespie simulation (individual life span) to calculate a fitness score given by a Lorentzian function centered the optimal phenotype. We also used the average protein number or the distribution of protein numbers over the individual life span as phenotypes; our results did not qualitatively change (Figure S12). We also changed the shape of the fitness function from a Lorentzian to a Gaussian or a step-like function with similar width; the results did not qualitatively change (Figure S12).

#### Mutations

Our simulations used a spherically symmetric 3D mutation scheme to permit co-variation in biophysical parameters in a single mutational step. The mutation step size was determined by the radius of the spherical mutation, which was a uniformly distributed random value between 0 and 1 (*r ∼ U*(0, 1)). Such a radial density produces a non-uniform density of mutations with highest densities close to the parental phenotype because volume scales as *r* ^3^. We tested homogeneous spherical mutation by substituting *r* in Eqs (5-7) with 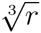 and a homogeneous cubic mutation where three uniformly distributed random value between −1 and 1 (*r_i_* ∼ *U* (−1, 1)) for each biophysical parameter. Both mutation schemes produced the same qualitative results (Figure S12). We also verified that mutating only one parameter at a time (1D mutation) and increasing the range of biophysical parameters to allow higher nonlinearity (10^−2^ ≤ *n_H_* ≤ 24) and weaker DNA dissociation constants (10^−2^ ≤ *K_D_* ≤ 10^3^) did not fundamentally change our results (data not shown).

#### Timescales of epimutation and stochastic gene expression

The rate of epimutation is sensitive to the frequency and magnitude of stochastic events. The magnitude of stochastic events is inversely proportional to the total number of molecules. Thus, we expect a higher rate of epimutation for smaller numbers of molecules. The rate of epimutation should also increase as the two modes become closer. Thus, we expect a higher rate of epimutation for larger α. Last, the protein degradation rate (γ) sets the timescale between stochastic events (i.e. faster protein degradation leads to more stochastic events per unit time during a Gillespie simulation). Thus, we expect a higher rate of epimutation for larger γ. In all tested cases, a higher rate of epimutation favored ES over GA (Figure S12).

#### Matching *α* and the ratio of optimal phenotypes

At high levels of nonlinearity, the lowest protein level is *k* · *α* and the highest protein level is *k*. A bistable, epigenetic switch has two solutions, each well-adapted to one of the environments only when the ratio 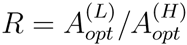 = *α* (Figure S1). Any mismatch between and *R* will disfavor epigenetic switching because an epimutation from an adapted mode will jump to a maladapted mode, after which the descendants must accumulate genetic mutations to further adapt. Although our simulations fixed *R* = *α* = 0.25, we verified that variable *α* evolved to ES with *α* ≈ *R* in fast fluctuating environments (high *ν*; Figure S13).

## ACKNOWLEDGMENTS

We thank Ryan Baugh, Kathleen Donohue, Sur Herrera Paredes, Katia Koelle, Mark Rausher, Joshua Socolar, and Marcy Uyenoyama for helpful feedback. This work was funded by a CONACYT graduate fellowship (MGS), the National Institutes of Health Directors New Innovator Award DP2 OD008654-01, and the Burroughs Wellcome Fund CASI Award BWF 1005769.01.

